# Subtype Heterogeneity and Epigenetic Convergence in Neuroendocrine Prostate Cancer

**DOI:** 10.1101/2020.09.13.291328

**Authors:** Paloma Cejas, Yingtian Xie, Alba Font-Tello, Klothilda Lim, Sudeepa Syamala, Xintao Qiu, Alok K. Tewari, Neel Shah, Holly M Nguyen, Radhika A. Patel, Lisha Brown, Ilsa Coleman, Wenzel M. Hackeng, Lodewijk Brosens, Koen M.A. Dreijerink, Leigh Ellis, Sarah Abou Alaiwi, Ji-Heui Seo, Mark Pomerantz, Alessandra Dall’Agnese, Jett Crowdis, Eliezer M. Van Allen, Joaquim Bellmunt, Colm Morrisey, Peter S. Nelson, James DeCaprio, Anna Farago, Nicholas Dyson, Benjamin Drapkin, X. Shirley Liu, Matthew Freedman, Michael C. Haffner, Eva Corey, Myles Brown, Henry W. Long

**Affiliations:** Department of Medical Oncology, Dana-Farber Cancer Institute, Brigham and Women’s Hospital, and Harvard Medical School, Boston, Massachusetts, USA; Center for Functional Cancer Epigenetics, Dana-Farber Cancer Institute, Boston, Massachusetts, USA; Translational Oncology Laboratory, Hospital La Paz Institute for Health Research (IdiPAZ) and CIBERONC, La Paz University Hospital, Madrid, Spain; Broad Institute of MIT and Harvard, Cambridge, MA, USA; Department of Urology, University of Washington, Seattle, WA, USA; Divisions of Human Biology and Clinical Research, Fred Hutchinson Cancer Research Center, Seattle, WA, USA; Department of Pathology, University Medical Center Utrecht, Utrecht University, Utrecht, The Netherlands; Department of Endocrinology, Amsterdam UMC, Amsterdam, The Netherlands; Department of Oncologic Pathology, Dana-Farber Cancer Institute and Harvard Medical School, Boston, MA, USA; Whitehead Institute for Biomedical Research, 455 Main Street, Cambridge, MA 02142, USA; Beth Israel Deaconess Medical Center and PSMAR-IMIM Lab. Harvard Medical School, Boston, Massachusetts, USA; Massachusetts General Hospital Cancer Center, Boston, Massachusetts, USA; Nancy B. and Jake L. Hamon Center for Therapeutic Oncology Research, Dallas, TX, USA; Harold C. Simmons Comprehensive Cancer Center, University of Texas Southwestern Medical Center, Dallas, TX, USA; Department of Internal Medicine, University of Texas Southwestern Medical Center, Dallas, TX, USA; Department of Data Science, Dana-Farber Cancer Institute, Harvard T.H. Chan School of Public Health, Boston, MA, USA; Division of Clinical Research, Fred Hutchinson Cancer Research Center, Seattle, WA, USA; Department of Pathology, University of Washington, Seattle, WA, USA

## Abstract

Neuroendocrine carcinomas (NEC) are tumors expressing markers of neuronal differentiation that can arise at different anatomic sites but have strong histological and clinical similarities. Here we report the chromatin landscapes of a range of human NECs and show convergence to the activation of a common epigenetic program. With a particular focus on treatment emergent neuroendocrine prostate cancer (NEPC), we analyzed cell lines, patient-derived xenograft (PDX) models and human clinical samples to show the existence of two distinct NEPC subtypes based on the expression of the neuronal transcription factors ASCL1 and NEUROD1. While in cell lines and PDX models these subtypes are mutually exclusive, single cell analysis of human clinical samples exhibit a more complex tumor structure with subtypes coexisting as separate sub-populations within the same tumor. These tumor sub-populations differ genetically and epigenetically contributing to intra- and inter-tumoral heterogeneity in human metastases. Overall our results provide a deeper understanding of the shared clinicopathological characteristics shown by NECs. Furthermore, the intratumoral heterogeneity of human NEPCs suggests the requirement of simultaneous targeting of coexisting tumor populations as a therapeutic strategy.

## Introduction

Neuroendocrine carcinomas (NEC) are high grade tumors, the most common being small-cell lung cancer (SCLC), that can also arise in the colon, prostate or the bladder among other anatomic sites. NECs are characterized by aggressive clinical behavior and a poor prognosis^1^. Histomorphologically, NEC comprise a heterogenous group of tumors which can have features of small cell carcinoma, and show expression of neuroendocrine markers including SYP, CHGA and INSM1^2^. Given these common characteristics, NECs constitute an unique clinicopathological entity despite their distinct anatomical origins^3^. From a genetic standpoint, NECs are often characterized by genomic aberrations in *RB1* and *TP53^4^*. The association of these genetic alterations with NEC etiology is exemplified in Merkel Cell Carcinoma (MCC). This aggressive NEC of the skin is caused in at least 60% of all MCC by clonal integration of Merkel cell polyomavirus DNA into the tumor genome with persistent expression of viral T antigens (MCCP; MCC Polyomavirus-positive). These viral proteins produce tumorigenesis by interfering with cellular tumor-suppressor proteins including RB1^5^. Alternatively, MCC can be caused by ultraviolet damage leading to highly mutated genomes causing a non-viral form of MCC (MCCN; MCC non-viral) that almost invariably carry mutations in *TP53* and *RB1^6^*. Despite these distinct etiologies, both forms of MCC have similar presentations, prognosis, and response to therapy. Gastrointestinal NEC (GINEC) is another relevant category of NECs that typically harbor *TP53* and *RB1* alterations and are clinically aggressive and highly proliferative. In that respect, the GINECs differ from the well differentiated GI carcinoids that, while also showing a NE phenotype, are typically clinically indolent^1^ and not associated with *TP53* and *RB1* alterations.

NEC can emerge either *de novo* or as a result of therapeutic pressure^7–10^. SCLC most often emerges *de novo* but can emerge after treatment of *EGFR* mutant NSCLC^11^. SCLC has been sub-classified based on the differential expression of the basic Helix-Loop-Helix (bHLH) transcription factors (TFs) ASCL1 and NEUROD1^12^. These neuronal lineage TFs have been implicated in lung development and in the maturation of resident neuroendocrine cells of the lung^13,14,15^. They are also involved in the carcinogenic process as shown in mouse models of SCLC where ASCL1 is required for tumor formation^16^.

Neuroendocrine prostate cancers (NEPC), in contrast, arises most frequently as a treatment emergent phenotype from prostatic adenocarcinomas after treatment designed to repress AR pathway activity^17^ and only rarely arises *de novo*. The improved potency and specificity of anti-AR therapy has led to an increased prevalence of NEPC^18^. NEPC has poor prognosis, very limited therapeutic options and is currently treated as a homogeneous disease. A better understanding of the molecular basis of NEPC is required in order to design more efficient targeted therapeutic options.

Here we utilize epigenetic profiling of a range of NECs to understand how the common phenotype is maintained across tumor types with particular focus on NEPC. We find that multiple bHLH transcription factors are implicated in a common epigenetic program across all NECs. Furthermore, epigenetic profiling reveals the existence of subtypes in treatment emergent NEPC in line with what has been described in *de novo* SCLC, despite arising from treated prostate adenocarcinoma. The subtypes coexist as separate sub-populations with distinct chromatin states within the same human NEPC specimens exhibiting heterogeneity that have clinical implications.

## Results

### Neuroendocrine carcinomas share a common landscape of DNA accessible regions

Histomorphologically, NECs show similarities that could result from activation of common transcriptional regulators^19^. To investigate the impact of chromatin accessibility in determining the NEC phenotype we profiled the epigenetic landscape of NECs arising in various anatomic locations using ATAC-seq and RNA-seq applied to PDX models of NEPC^20^, SCLC^20,21^, and Merkel cell carcinoma (MCC) as well as GINEC clinical samples (Table 1). As NEC can emerge from a preexisting adenocarcinoma (AD), as typified by NEPC^22^ and occasionally by SCLC^18,23^, we hypothesized that those histologies are extremes of a spectrum of tumor progression. To determine how the chromatin state differs between neuroendocrine (NE) and adenocarcinoma (AD) by ATAC-seq analysis, we also generated data from metastatic prostate adenocarcinoma PDX models and used TCGA data for primary Prostate ADs (PRAD) and non-small cell lung adenocarcinomas (LUAD)^24^. We obtained high quality data with the fraction of reads in peaks (FrIP) scores in the range of 10-35 and peak numbers in the range of 25-75k (Table 1). Replicate profiling of two samples showed high concordance (Suppl. Figure 1a). Unsupervised principal-component analysis (PCA) performed on the ATAC-seq data revealed that the NECs cluster together indicating a convergent chromatin state, in contrast to the ADs that are segregated by anatomic site (Figure 1a). The sample-sample correlation of the ATAC-seq peaks also supports the result that NECs are more similar to each other than to their AD counterparts from the same tissue (Figure 1b), although the MCCs are particularly homogeneous (one MCCN is tightly clustered with five MCCPs). These analyses also clustered prostate PDXs and primary human tumors together emphasizing, in terms of the chromatin state, the value of the prostate LuCaP^25,26^ PDXs to model human prostate cancer, as previously validated by histological and molecular characterization^25,26^.

**Figure 1.**
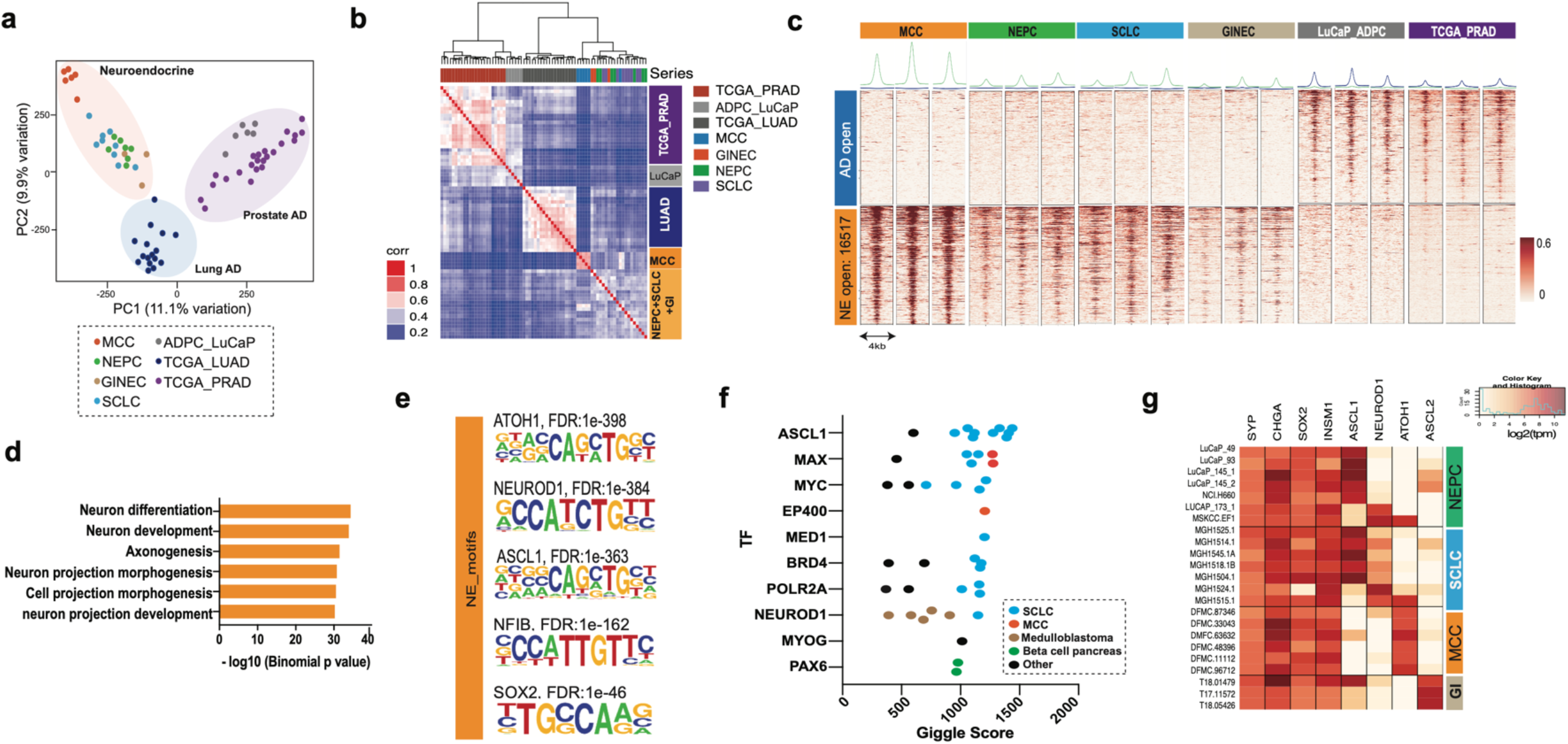
NE carcinomas share a common chromatin state independent of their anatomical origin. **(a)** PCA analysis of ATAC-seq data of NECs including Merkel Cell Carcinoma (MCC), Neuroendocrine prostate cancer (NEPC), Neuroendocrine gastrointestinal (GINE) and Small cell lung cancer (SCLC). The plot also includes Prostate adenocarcinoma (PDX models and TCGA primary tissues) and Lung adenocarcinoma (TCGA primary tissues). (**b**) Hierarchical clustering of the pairwise Pearson’s correlation of the ATAC-seq signal across the distinct tumor types. (**c**) Heatmap representation of the differential regions between ADs and NECs. Each row is a peak location and each column is a sample. Shown above each column are the composite tag density plots for the AD sites (blue) and NE sites (green). (**d**) Gene Ontology pathways enriched in genes with nearby NE-specific accessible regions shown in (c). (**e**) Top results from motif analysis of the NE-specific accessible regions. (**f**) Public ChIP-seq datasets showing the highest overlap with the NE-specific accessible regions annotated by tissue type as determined by CistromeDB toolkit. The TFs are ordered by the top scoring dataset of each type. (**g**) Expression of NE markers and bHLH TFs across all the NEC samples in our study displayed as a heatmap.

To investigate epigenetic drivers involved in the NE chromatin state, we performed a supervised analysis of the DNA accessibility between ADs and NECs and found a high number (n:16517, Padj<0.001, log2(FC) >2) of NE-specific accessible sites shared across all NE tumor types (Figure 1c). This represents a major difference in chromatin organization that we further investigated by GREAT analysis^27^ that associates genomic regions with nearby genes and then examines the enrichment of Gene Ontology (GO) pathways. Genes near NE-specific DNA accessible regions showed a significant enrichment in pathways for neural differentiation, development, morphology and axogenesis (Figure 1d). Next, we used HOMER^28^ to investigate NE-specific sites for enrichment of TF DNA-binding motifs. This analysis revealed significant enrichment for motifs of the basic Helix-Loop-Helix (bHLH) TF family, specifically for ATOH1, ASCL1 and NEUROD1 as well as motifs for NFIB, SOX2 and NKX2-1 (Figure 1e and Table 2). ATOH1 has been implicated as a lineage transcription factor in MCC^29,30^, while ASCL1 and NEUROD1 have been suggested to have a corresponding role in SCLC^16,31^. NFIB is a TF previously implicated in rewiring the chromatin structure in SCLC^32^, while SOX2 and NKX2-1 are also known to be associated with SCLC^33,34^. Examining what motifs co-occur in the ATAC-seq peaks we observed that the module of the main three motifs (ASCL1, ATOH1 and NEUROD1) occurs very frequently combined with either SOX2 or NFIB or with both of them simultaneously (Suppl. Figure 1b).

By comparing the NE-specific sites to published ChIP-seq profiles compiled in CistromeDB^35^, we identified TFs whose published binding sites have the highest overlap with the NE-specific ATAC-seq peaks as quantified by GIGGLE score^36^. Consistent with the observed shared epigenetic program amongst NECs of different tissue origins, the top overlapping ChIP-seq datasets were generated from SCLC, MCC or neural lineages (Figure 1f). In particular, binding profiles of ASCL1 (SCLC), NEUROD1 (SCLC and medulloblastoma), and MAX (MCC)^5^ had the highest overlap scores with NE-specific accessible chromatin(Figure 1f). We next analyzed the expression of these TFs and other NEC-associated factors within our study samples. As expected, we observed a strong commonality in the expression of NE markers (*SYP, CHGA and INSM1*) and the stemness TF, *SOX2* (Figure 1g) across NECs. We also observed a more mutually exclusive expression pattern of bHLH TFs including *ASCL1* or *NEUROD1* in both NEPC and SCLC, *ASCL1* and *ASCL2* expression in GI-NECs and *ATOH1* expression in MCC (Figure 1g). This suggests tumor and organ specific TF drivers of this common neuroendocrine epigenetic state. Overall our results across a diverse panel of NECs show a common landscape of DNA accessible regions with expression of varying TFs within the same family.

### Treatment emergent NEPC can be subclassified based on the expression of ASCL1 and NEUROD1

To explore heterogeneity in the TF regulation of the NEC epigenetic state, we performed an unsupervised analysis of the ATAC-seq data restricted to the NECs. Regardless of tissue of origin, NECs expressing *ASCL1* and/or *ASCL2* were tightly clustered together and were separate from NECs expressing *ATOH1* or *NEUROD1* (Suppl. Fig 2a). Furthermore, the similarity in terms of the DNA accessibility shown by SCLC and NEPC depends on the status of *ASCL1* or *NEUROD1* expression but not on the tumor type (Suppl. Fig. 2b). An unsupervised analysis of the DNA accessibility in just prostate samples (Figure 2a) showed clear grouping associated with expression of *AR* (all adenocarcinomas), *ASCL1* or *NEUROD1* with the same clustering being apparent by analysis of RNA-seq data of those same prostate samples (Suppl. Fig. 2c). Although neuroendocrine subtypes based on the expression of those TFs have been previously described in SCLC^12,16^, the existence of these subtypes in treatment emergent NEPC was unanticipated as ASCL1 and NEUROD1 have been specifically associated with lung neuroendocrine cells^13,14,15^, the putative cell of origin of the *de novo* SCLC.

**Figure 2.**
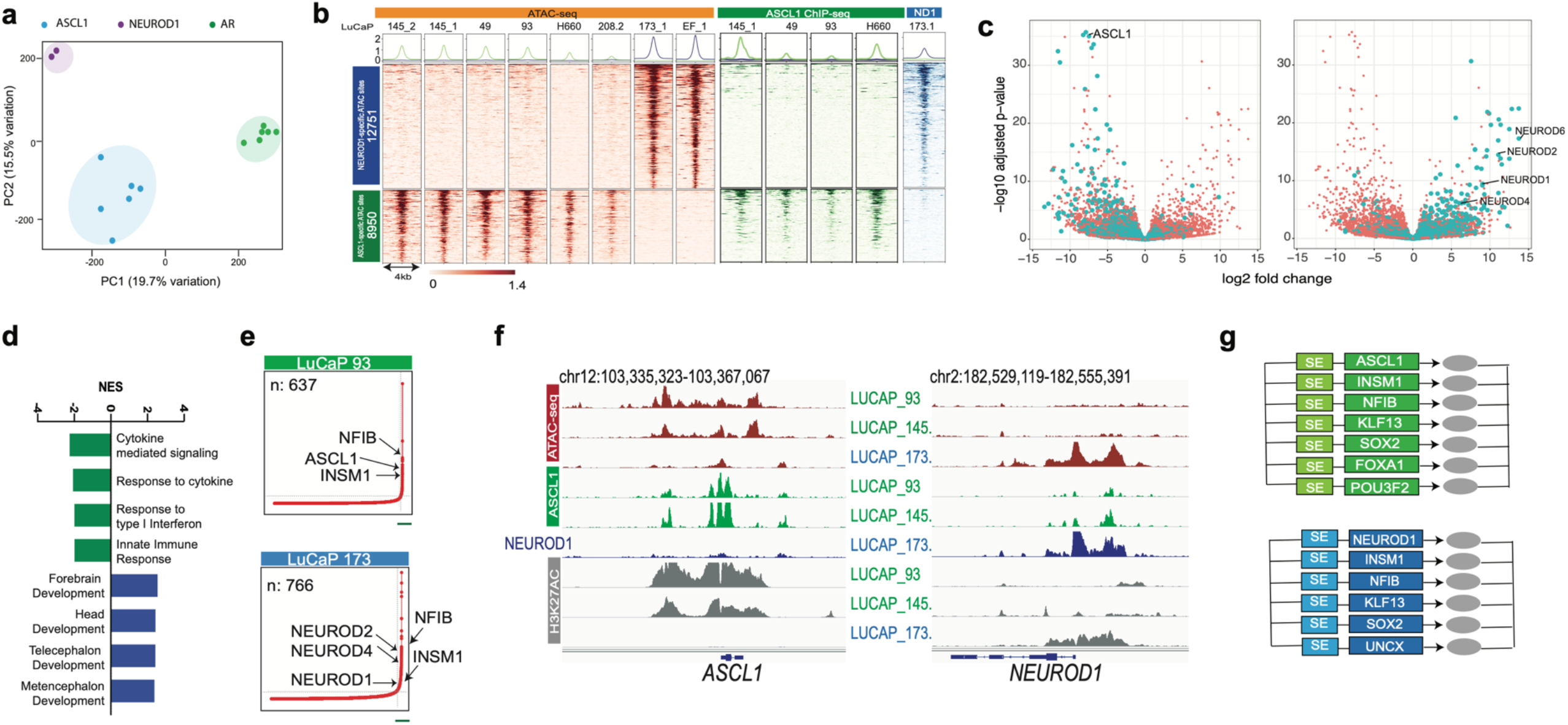
NEPC shows tumor subtypes based on the differential expression of the transcription factors ASCL1 and NEUROD1. **(a)** PCA analysis of NEPC and ADPC PDXs ATAC-seq data. Samples are color coded by the dominant TF expressed in that sample. (**b**) Heatmap representing the differential ATAC-seq regions between NEPC subtypes. ASCL1 ChIP-seq (green) and NEUROD1 ChIP-seq (blue) are shown for the indicated samples at the same sites. (**c**) Association between differential ATAC accessible sites and differential gene expression. Each volcano plot depicts RNA-seq log2 fold change and adjusted p-value calculated by DESeq2. Each dot represents one gene: green indicates a differential ATAC peak is with 50kB and orange indicates no such peak. ASCL1-specific ATAC accessible regions and genes upregulated in ASCL1 subtype (left) or NEUROD1-specific accessible regions and genes upregulated in NEUROD1 subtype (right). (**d**) GSEA pathway analysis enriched in ASCL1 subtype (green) and NEUROD1 subtype (blue). (**e**) Superenhancer analysis of a representative case of the ASCL1 subtype (top) and NEUROD1 subtype (bottom). (**f**) Representative IGV tracks at the *ASCL1* and *NEUROD1* gene. ATAC-seq tracks are in green, ASCL1 ChIP-seq in green, NEUROD1 ChIP-seq in blue and H3K27ac in gray. The loci are marked by subtype specific SEs with preferential binding of their respective TF. (**g**) Circuits of lineage transcription factors specific for the ASCL1 subtype (green) and NEUROD1 subtype (blue).

Next we aimed to identify the differential DNA accessibility associated with the ASCL1 and the NEUROD1 NEPC subtypes. Supervised analysis comparing *ASCL1* and *NEUROD1* expressing NEPC samples identified 2863 ASCL1- and 4873 NEUROD1-specific accessible regions (FDR<0.01, log2(FC) > 2) (Figure 2b). We next interrogated the NEPC subtype-specific sites in SCLC and observed a similar patterns of chromatin accessibility at these TF-specific genomic regions in SCLC samples that, in addition, displayed an association between the chromatin state and the differential expression of *ASCL1* and *NEUROD1* (Suppl. Fig. 2d). Notably, SCLC cases that coexpress *ASCL1* and *NEUROD1* showed combined accessibility at the two sets of regions (Suppl. Fig. 2d). This result underlines the striking similarity in the chromatin state of the tumor subtypes both in SCLC and NEPC. It is important to note that despite the clear differences in accessibility associated to the subtypes, still the large majority of open chromatin sites are shared between these two subtypes as expected given the NE characteristics in common for both subtypes (Suppl. Fig. 2e).

To further characterize the chromatin differences between the subtypes we investigated the relationship between TF-subtype specific chromatin accessibility and the ASCL1 and NEUROD1 genomic binding. To that aim, we performed ChIP-seq analysis for each of the two TFs in NEPC models that expressed *ASCL1* or *NEUROD1* (Table 3). This analysis identified thousands of highly conserved binding sites with both overlapping and differential sites for each of the two TFs (Suppl. Fig. 2f) that were enriched for the expected consensus motifs for ASCL1 and NEUROD1 respectively (Suppl. Fig 2g). Importantly, TF-specific regions of differential chromatin accessibility were bound by the corresponding TF, but not the other (Figure 2b). This result is consistent with a role for ASCL1 and NEUROD1 in maintaining the chromatin state in their respective subtypes.

We next sought to confirm that the enhancers specific to the ASCL1 and NEUROD1 subtypes are associated with expression of nearby genes by using the expression data generated from the same samples (Figure 2c). As expected, differential expression analysis showed that *ASCL1* was one of the most upregulated genes in the ASCL1 set, while conversely, the NEUROD1 set showed upregulation of several NEUROD family members including *NEUROD1/2/4/6* (Figure 2c). Consistent with these differentially accessible regions being functional, we observed a substantial association between differential DNA accessibility and differential gene expression (Figure 2c). Gene Set Enrichment Analysis (GSEA) to identify pathways differentially over-represented in each of the two subtypes showed that ASCL1 associated gene expression was enriched in GO pathways of response to cytokines^37^ while the NEUROD1 associated expression was enriched in brain development pathways (Figure 2d).

The binding of ASCL1 and NEUROD1 TFs to their own promoters and nearby enhancers suggests they are working as lineage TFs (LTFs) in NEPC. LTFs are known to auto-activate their own expression by binding to super-enhancers (SE) establishing a positive feedback loop. In addition, LTFs form circuits of core TFs driven by the activation of super-enhancers (SE) promoting the transcriptional program required to maintain the lineage^38^. Both ASCL1 and NEUROD1 are known to be lineage transcriptional factors in neuronal systems^39,40^. To investigate a potential LTF behavior of both TFs in NEPC, we performed SE analysis by H3K27ac profiling of ASCL1 and NEUROD1 NEPCs and found that all models showed SE activation in common at a number of TFs (*INSM1* and *NFIB*) regardless of the tumor subtype. In addition, we found differential SEs at either *ASCL1* or *NEUROD1* (and other family members) in accordance with their expression status (Figure 2e). Based on those characteristics both ASCL1 and NEUROD1 can be considered as LTFs in NEPC with binding to SEs and activation of their own expression (Figure 2e, 2f). We next identified the core circuit of TFs associated with each of the two subtypes applying a previously described method to identify interconnected auto-regulated loops^38^. We identified distinct but highly overlapping sets of TF circuits in these two subtypes (Figure 2g).

Taken together, our results provide clear evidence of the existence of two molecular subtypes in NEPC model systems. These subtypes share NE phenotypic characteristics but differ in the expression of *ASCL1* and *NEUROD1*, which is associated with distinct chromatin landscapes and gene expression profiles.

### Analysis of tumor heterogeneity in NEPC liver metastases

Next we aimed to determine if the results from the model systems can be extended to human clinical NEPC. First we interrogated expression levels of *ASCL1* and *NEUROD1* in tumor tissues from two cohorts of NEPC metastases^22,25^. In contrast to the mutually exclusive expression of the two TFs that we observed in NEPC PDXs (Figure 1g), clinical samples showed a range of coexpression. The *ASCL1* expression was higher in the majority of the metastases accompanied by a lower and more variable expression of *NEUROD1* for almost all the cases (Figure 3a, Suppl. Fig 3a).

**Figure 3.**
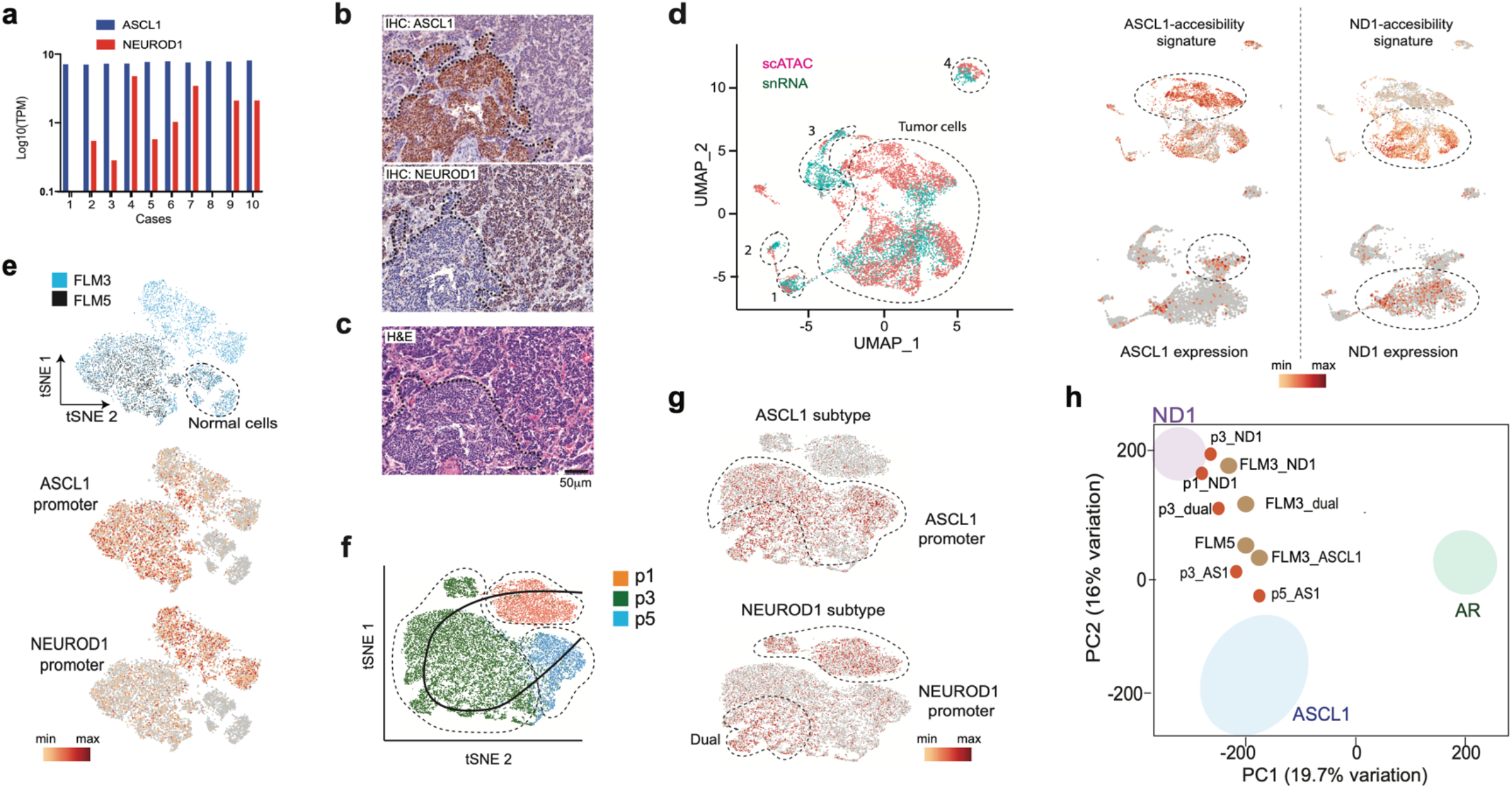
Single cell analysis reveals that NEPC sub-types co-exist in human metastasis and contribute to inter- and intra-tumoral heterogeneity. (**a**) Plot of *ASCL1* and *NEUROD1* expression in NEPC tissues from a clinical cohort (Labreque *et al*. 2019)^25^. TPM: Transcripts per million. (**b**) Immunohistochemical analysis of FLM3 (ASCL1 staining in top panel and NEUROD1 staining in the middle panel) showing intratumor heterogeneity. (**c**) Hematoxylin and eosin staining illustrating the distinct histologies for the two subpopulations. (**d**) Combined analysis of the scATAC-seq and snRNA-seq in FLM3 using SEURAT (left). Markers specific for normal cell populations enabled assignment of clusters: 1, vascular cells; 2, stromal cells; 3, hepatic cells; 4, macrophages. Accessibility at the top 30 differential ATAC-seq regions between ASCL1 and NEUROD1 subtypes identified by bulk analysis (top right). Analysis of *ASCL1* and *NEUROD1* expression in the snRNA-seq analysis (bottom right). This analysis matches cells with TF expression and the corresponding differential DNA accessibility for each subtype. (**e**) t-SNE analysis of the combined FLM3 (blue) and FLM5 (black) scATAC-seq data (top). The lower two plots show the differential accessibility at *ASCL1* promoter (middle) and *NEUROD1* promoter (bottom). (**f**) t-SNE plot of the combined PDX data showing the projected trajectory across all of the cells in the PDX analysis. (**g**) t-SNE plot of the combined PDX showing the differential accessibility of the combined PDX dataset at the *ASCL1* promoter indicating the ASCL1 subtype (top) and *NEUROD1* promoter showing the NEUROD1 subtype (bottom) the results also show a cluster with dual accessibility at *ASCL1* and *NEUROD1* promoters. (**h**) Projection of the scATAC-seq clusters for FLM3 and 5 (light brown dots) and PDX passes (orange dots) within the PCA space defined in Figure 2a.

To investigate whether these TFs are co-expressed in the same tumor cells or in distinct tumor sub-populations, we studied five distinct fragments of liver metastasis (FLM) obtained at autopsy from a patient diagnosed with NEPC available as both OCT frozen and formalin-fixed paraffin-embedded material (Suppl. Fig 3b) and performed RNA-seq to assess expression levels of *ASCL1* and *NEUROD1*. We observed a range of coexpression of the two TFs with FLM3 showing the highest relative expression of *NEUROD1* to *ASCL1* (Suppl. Fig 3c). We next performed immunohistochemical (IHC) analysis for ASCL1 and NEUROD1 protein expression on FLM3. The staining showed clear intratumoral heterogeneity in terms of ASCL1 and NEUROD1 staining that defined two separated tumor populations (Figure 3b). Correlation with histomorphological features showed that the two distinct cell populations also differ in their histological characteristics. ASCL1 positive cells had a sheet-like growth pattern and spindle cell morphology, whereas NEUROD1 positive cells appeared to grow in smaller cell clusters with pronounced nuclear molding and focal pleomorphic giant cells (Figure 3c). We next performed double staining of ASCL1 and NEUROD1 by immunofluorescence in FLM3 to investigate potential co-expression in tumor cells and observed that the vast majority of the cells showed an anticorrelated expression of the two TFs (Suppl. Fig 3d). We extended this analysis by investigation of two additional NEPCs from different patients by IHC and also observed the existence of this type of intratumor heterogeneity (Suppl. Fig 3e).

We next investigated the two observed intra-tumoral populations by single cell chromatin (scATAC-seq) and expression (snRNA-seq) analysis. We selected FLM3 that showed the highest NEUROD1 expression and FLM5 that had the lowest, almost 200-fold lower than *ASCL1* (Suppl. Fig 3c). We isolated nuclei from frozen sections of FLM3 and performed scATAC-seq and snRNA-seq to assign the *ASCL1* and *NEUROD1* expression with the corresponding chromatin state. The unsupervised tSNE clustering of the scATAC-seq resulted in multiple clusters that we analyzed for differential accessibility at *SOX2* promoter to distinguish tumor and normal cells. Based on accessibility to *SOX2* the fraction of the tumor cells represented around 80% (Supp. Fig. 3f). Notably, we could distinguish the clusters that correspond to the two tumor subtypes based on the differential accessibility to the *ASCL1* and *NEUROD1* promoters (Suppl. Fig. 3f). The ASCL1 and NEUROD1 clusters also show differential accessibility at the top ATAC differential regions identified by bulk analysis (Suppl. Fig. 3g). A small number of cells displayed chromatin accessibility at both differential regions and the *ASCL1* and *NEUROD1* promoters, that we labelled as “dual” (Suppl. Fig. 3f,g). We next analyzed the snRNA-seq to identify the tumor cells that express *ASCL1* and *NEUROD1* and then integrated this dataset with the scATAC-seq using SEURAT^41^ (Figure 3d). This integration enabled the assignment of normal cells based on the expression of specific markers. Crucially, we observed that cells with either the ASCL1 or NEUROD1 accessibility signature developed from bulk data preferentially express the corresponding TF (Figure 3d). Thus, the integrated single cell analysis clearly shows that the ASCL1 and NEUROD1 sub-types exist as separate subpopulations possessing similar epigenetic features as in their respective model systems.

We next investigated the FLM5 sample, which has the highest expression of *ASCL1* by scATAC-seq analysis. In accordance with the RNA-seq, the t-SNE analysis showed a single cluster of the FLM5 tumor cells with accessibility at *ASCL1* promoter but not at *NEUROD1* (Suppl. Fig. 3h). The integrated scATAC analysis of FLM3 and FLM5 revealed that 99% of the FLM5 tumor cells overlap the FLM3 ASCL1 cluster (Figure 3e) indicating that those cells have identical chromatin accessibility.

The identification of the two coexisting subtypes in NEPC metastases motivated the further investigation in the PDX model derived from this patient (Suppl. Fig 3b). This PDX model was characterized by RNA-seq analysis of multiple samples from passage 1 (p1) up to p17. The system showed dramatic changes in the relative expression of *ASCL1* and *NEUROD1* across passages, evolving from high *NEUROD1* expression and barely detectable expression of *ASCL1* at p1, to a relative co-expression at p3, to high *ASCL1* expression and extremely low *NEUROD1* expression from p5 onwards (Suppl. Fig 3i). We selected samples from p1, p3 and p5 for analysis by scATAC-seq analysis and performed the trajectory inference of the tumor cell evolution by analysis of the scATAC-seq results using Monocle^42^ (Figure 3f). The tSNE analysis of the scATAC-seq data revealed homogenous populations at p1 and p5 consistent with the NEUROD1 and ASCL1 sub-types respectively, which was also validated by IHC (Suppl. Fig 3j), while p3 showed coexistence of the sub-type chromatin accessibility patterns (Figure 3f and 3g). Therefore, the inferred trajectory ordered the tumor cells along a continuum based on DNA accessibility with the pure ASCL1 and NEUROD1 tumor populations as the two edges of the trajectory (Figure 3f). Plotting the aggregated scATAC-seq by TF cluster from FLMs and PDXs in the PCA space defined by the model systems in Figure 2a further illustrates the PDX changes from NEUROD1 to ASCL1 while also validating the NEPC model systems we have used (Figure 3h). All together, these results demonstrate subtype heterogeneity in human NEPC metastases and that these subtypes show distinct epigenetic characteristics and a divergent dynamic behavior.

### The NEPC sub-types are distinct but related clones in the patient metastasis

We next sought to investigate the genetic characteristics of the NEC samples using whole exome sequencing (WES) and copy number variation (CNV) inferred from the ATAC-seq data^43^. Inference of *RB1* genetic status from the bulk ATAC-seq data showed biallelic loss in all the NEPC PDX models but not in the ADPCs (Suppl. Fig. 4a) as previously reported for these NEPC models^20^. The same approach was applied genome-wide to the scATAC-seq clusters identified by the tSNE analysis on FLM3 and FLM5. The results show an overall similarity in the CNVs across these clusters (Figure 4a). For instance, we observed heterozygous losses in all of chr16 and parts of chr2 and chr13 in both ASCL1 and NEUROD1 clusters. In addition, we found a focal heterozygous loss at *PTEN* on chr10 in both clusters. However, clear CNV differences existed, including a 20MB amplification on chr14p and a chr7p amplification that are only present in the NEUROD1 cluster. Notably, CNVs of the ASCL1 component in FLM3 showed almost identical characteristics with the ASCL1 cluster in FLM5 (Pearson correlation = 0.97) while showing lower correlation to the NEUROD1 cluster within the same fragment (Pearson correlation = 0.81) (Figure 4b). These same CNV alterations along with others were observed in PDX passages. Specifically, p1 and a p3 cluster had CNVs that were most similar to the NEUROD1 profile, while the other p3 cluster and the p5 were most similar to the ASLC1 profile (Figure 4c, Suppl. Fig. 4b). This result supports the hypothesis that the dynamic changes in the PDX model are due to the preferential outgrowth of the ASCL1 clone that started at a very low fraction in the initial passage. WES analysis of the FLM samples, though derived from bulk tissue, validated the scATAC-seq inferred CNV alterations including amplifications on chr14p and chr7p (Suppl. Fig 4c).

**Figure 4.**
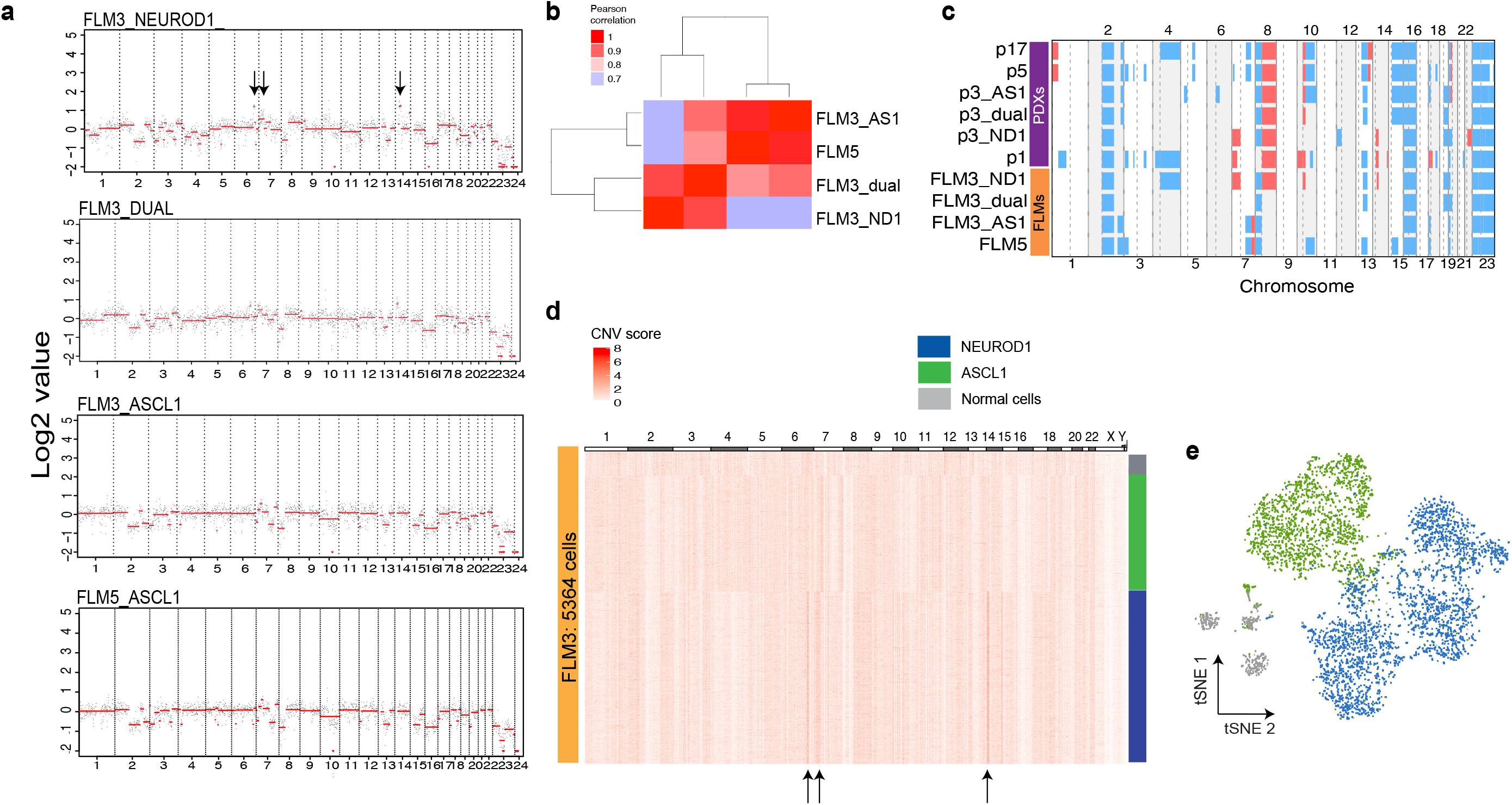
The NEPC sub-types are distinct clones. **(a)** Genome wide CNV profiles inferred from the scATAC-seq clusters in FLM3 and FLM5. Black dots are values in 1MB regions and the red line is the result of running a segmentation algorithm on the data (see Methods). Arrows point to differences seen in CNVs across the clusters. (**b**) Sample pairwise Pearson’s correlation of the CNV profiles. (**c**) Summary heatmap of the scATAC-seq inferred CNV alterations across all of the patient samples and PDX models profiled (blue represents losses and red represents gains). (**d)** Heatmap of the single cell CNV analysis of FLM3 where each column is a 2MB bin tiled across the genome and the rows are individual cells that have been clustered with K-means. Arrows point to CNV differences observed here and in the cluster level analysis. (**e**) tSNE plot of FLM3 scATAC-seq data colored by the cluster each cell was partioned into by the inferred CNV alterations. Those three clusters clearly correspond to NEUROD1 (blue), ASCL1 (green) and normal cells (gray).

Finally, we extended the CNV analysis of FLM3 to the single cell level following the method of Satpathy et al.^43,44^. The K-means clustering of the cells based on the CNVs distinguished normal cells with no alterations from two additional clusters, one with the chr7p and chr14p amplifications corresponding to the NEUROD1 tumor, and one without these alterations corresponding to the ASCL1 tumor (Figure 4d). We next marked the identity of the cells from the three clusters defined in the genetic analysis within the scATAC-seq tSNE plot from this sample and showed a strong correspondence to the groupings defined by the epigenetic analysis (Figure 4e). Importantly, this single cell resolution enabled a more refined interpretation of the dual accessibility initially identified in the analysis of the scATAC-seq results. The CNV inference of that cluster showed that it is composed of a mix of cells belonging to either ASCL1 or NEUROD1 clones (Suppl. Fig. 4d) and is not an independent clone. All together our results show the existence of distinct genetic clones associated with each of the two NEPC epigenetic subtypes in this patient, likely derived from a common ancestor given their substantial CNV profile overlap.

## Discussion

Poorly differentiated NECs are a class of high-grade tumors that arise at different anatomical sites and typically express markers of neuroendocrine differentiation (CHGA, NCAM1, and SYP). Our results build considerably on previous work with RNA-seq and cell lines^19,45^ and provide a molecular rationale for the shared histopathological behavior of these tumors based on a common epigenetic state regardless of anatomic origins or the distinct tumor-initiation mechanisms. This epigenetic convergence is associated with the expression of bHLH TFs known for specifying neural lineages. Notably, distinct members of the bHLH family are expressed by the different NECs, suggesting that a variety of TFs can maintain the NE fate.

The similarity in the chromatin state across NECs is particularly pronounced between NEPC and SCLC, which is surprising given the distinct cells of origin in these neoplasms. Although SCLC can arise as a treatment emergent phenotype, it mostly arises *de novo* and is thought to originate from resident neuroendocrine cells of the lung known to express either ASCL1 or NEUROD1^13,14,15^. In contrast, NEPC is primarily observed as a treatment emergent tumor that arises through a process of transdifferentiation from a preexisting adenocarcinoma^22^. Thus, despite the initial adenocarcinomas having a radically different chromatin state, the pattern of DNA accessibility observed in the resulting NEPC becomes almost indistinguishable from that in SCLC. More unexpected was our observation that treatment emergent NEPC also shows subtypes based on the expression of *ASCL1* and *NEUROD1* as in SCLC, since those TFs have been previously associated with lung development. Despite both subtypes having the NE phenotype, with expression of *SYP*, *CHGA* and *INSM1* and exhibiting largely similar chromatin states, they still differ in the activation of specific sets of enhancer sites and transcriptional programs.

Importantly, we show a fundamental difference between the representation of the subtypes in the PDX models as compared to human clinical samples. In contrast to the mutually exclusive expression of *ASCL1* and *NEUROD1* in model systems, tissues from NEPC clinical cohorts show coexpression of *ASCL1* and *NEUROD1* at varying levels. Single cell analyses of a set of metastatic samples from the same patient revealed the presence of two distinct tumor populations that coexist within the metastasis. This observation emphasizes that PDXs, despite being good models of the human disease, still offer limitations to illustrate the complexity observed in primary tissues. In fact, those limitations could have precluded a better characterization of subtype coexistence in SCLC^46^ that has mainly been described as homogeneous subtypes^12^. Our results show the existence of subtypes in clinical samples of NEPC and demonstrate heterogeneity in terms of the chromatin state.

CNV inference from scATAC-seq clusters demonstrated the existence of distinct clones associated with the two NEPC epigenetic subtypes. Based on our results we propose that the ASCL1 clone originated by transdifferentiation from an adenocarcinoma cell and there was subsequently subclonal evolution to the NEUROD1 state (Suppl. Fig 4e). This is supported by recent experiments that show tumors in Myc-driven SCLC GEMMs that start as pure *ASCL1* can develop *NEUROD1* expression both *in vitro* and *in vivo^46^*. Both NEPC clones then seed the liver metastases with the earlier ASCL1 clone being more common. It has been observed in CRPC studies that metastasis-to-metastasis seeding can occur and that may be contributing here as well^47^. However, we cannot rule out an alternative hypothesis where separate transdifferentiation events occur for the ASCL1 and NEUROD1 clones from the preexisting adenocarcinoma.

The intra-tumor heterogeneity in NEPC that we have observed here has direct clinical implications. In SCLC these subtypes have been linked to specific drug targets or predictors of drug response. For example, DLL3 is a target for an antibody-drug conjugate in the ASCL1 subtype^48,49^ and AURKA is a target for small molecule inhibitors such as alisertib in the NEUROD1 subtype^50^. With this new understanding of sub-type heterogeneity based on NEUROD1 and ASCL1 in NEPC, therapeutic strategies that target one but not the other will rapidly succumb to outgrowth of the resistant subpopulation. Altogether, our results illustrate the epigenetic complexity that exists in clinical tumors and provide a rational basis for targeting the inter and intra tumoral heterogeneity as a therapeutic strategy in NEPC.

## ACKNOWLEDGEMENTS

P.C. acknowledges funding from the Ministry of Economy and Competitiveness, Instituto de Salud Carlos III (Institute of Health Carlos III)—PI18-01604. H.W.L., M.B., E.C., P.S.N. acknowledge support from NIH grant P01 CA163227-06A1. C.M. & P.S.N. acknowledge P50 CA097186-16A1, while P.S.N. also acknowledges support from W81XWH-17-1-0415. N.J.D, B.J.D and A.F.F are supported by U01CA220323 (N.J.D).

## Methods

### Clinical samples

Tissue samples were collected within 8 hours of death from patients who died of metastatic CRPC. All patients signed informed consent for a rapid autopsy, under the aegis of the Prostate Cancer Donor Program at the University of Washington. Hematoxylin and eosin-stained slides from each case were reviewed by a pathologist to confirm the presence of tumor cells. The Institutional Review Board of the University of Washington (IRB #2341) approved this study.

### Nuclei preparation

Fragments of frozen tissues (PDX models) or 50 um sections (liver metastases) were cut and resuspended in 300 ul of cold 3-detergent-ATAC-Resuspension Buffer (RSB) containing 0.1% NP40, 0.1% Tween-20, and 0.01% Digitonin. Tissues were dounced 10 times each with a loose and a tight pestle each until homogenization was complete. The homogenate was then transferred to a 1.5 ml pre-chilled microfuge tube and incubated on ice for 10 minutes. After lysis, 300 ul of ATAC-RSB containing 0.1% Tween-20 was added and the tubes were inverted to mix. Lysates were filtered through a 40 um cell strainer and nuclei were centrifuged for 10 minutes at 1500 RCF in a pre-chilled (4 °C) fixed-angle centrifuge. Nuclei were resuspended with 300 ul of ATAC-RSB containing 0.1% Tween-20 and counted with a hemocytometer using Trypan blue stain.

### ATAC-seq

100,000 nuclei were resuspended in 50 ul of transposition mix (25 ul 2x TD buffer, 2.5 ul transposase (100 nM final), 16.5 ul PBS, 0.5 ul 1% Digitonin, 0.5 ul 10% Tween-20, 5 ul H2O]^52^. Transposition reactions were incubated at 37 °C and shaken at 1000 RPM for 30 minutes on a thermomixer. Transposed DNA was purified using Qiagen columns. Libraries were amplified as described previously^53^. 35-bp paired-end reads were sequenced on a NextSeq instrument (Illumina).

### ChIP-sequencing

Nuclei isolated as previously described were crosslinked with 1% Formaldehyde for 10 minutes for H3K27Ac ChIP-seq. For ASCL1 and NEUROD1 ChIP-seq nuclei were crosslinked in 2 steps with 2 mM of DSG (Pierce) for 45 minutes at room temperature followed by 1 ml of 1% Formaldehyde for 10 minutes. Crosslinked nuclei were then quenched with 0.125 M glycine for 5 minutes at room temperature and washed with PBS. After fixation, pellets were resuspended in 500 ul of 1% SDS (50 mM Tris-HCl pH 8, 10 mM EDTA) and sonicated for 5 (H3K27ac) or 10 (ASCL1 and NEUROD1) minutes using a Covaris E220 instrument (setting: 140 peak incident power, 5% duty factor and 200 cycles per burst) in 1 ml AFA fiber millitubes. Chromatin was immunoprecipitated with 10 ug of H3K27Ac antibody (Diagenode cat# C15410196), 10 ug of ASCL1 antibody (abcam ab74065) or 10 ug of NEUROD1 antibody (Cell Signaling mAb #4373). 5 ug of chromatin was used for H3K27Ac ChIPs, and 40 ug of chromatin was used for ASCL1 or NEUROD1 ChIPs. ChIP-seq libraries were made using Rubicon kit and purified. 75-bp single-end reads were sequenced on a Nextseq instrument (Illumina).

### Single nuclei ATAC-seq and RNA-seq

Nuclei were prepared as described previously. For scATAC-seq nuclei were transposed according to the OMNI-ATAC protocol^52^. ~7,000 cells were targeted for each sample and processed according to the 10x Genomics scATAC-seq sample preparation protocol (Chromium Single Cell ATAC Library & Gel Bead Kit, 10x Genomics). For snRNA-seq, nuclei prepared the same way were used directly in the 10x Genomics snRNA-seq protocol (Chromium Single Cell 3’ v2 Reagent Kit, 10xGenomics).

### RNA-seq

A fragment of frozen tissues (PDX models) or 50 um sections (liver metastases) were cut and homogenized in 1 ml of AllPrep DNA/RNA Mini Kit (Qiagen) using a plastic pestle (Cole-Palmer #44468-23). DNA and RNA were simultaneously isolated. 500 ng RNA was used to prepare libraries using the NEBNext. Ultra™ RNA Library Prep Kit for Illumina. RNA quantity and quality were assessed on an Agilent 2100 Bioanalyzer. For all RNA-seq, reads were sequenced on a NextSeq 500 instrument (Illumina).

### Whole exome sequencing

DNA extraction on frozen PDXs, human FLMs and adjacent normal tissue was performed using the AllPrep DNA/RNA Mini Kit (Qiagen). Whole exome sequencing was performed by Novogene using their standard protocols. Briefly, 1000 ng of genomic DNA were used as input to generate sequencing libraries using the Agilent SureSelect Human All Exon Kit. Captured libraries were enriched by PCR, purified, quantified using the Agilent Bioanalyzer 2100 system, and subsequently sequenced using the NextSeq 500 instrument (Illumina).

### IHC

Immunohistochemical and immunofluorescence studies using ASCL1 (clone 24B72D11.1, cat. no. 556604, BD biosciences, San Jose, CA) and NEUROD1 (clone EPR17084, cat. no. ab205300, Abcam, Cambridge, MA) specific antibodies were carried out on archival formalin fixed paraffin embedded tissues. In brief, 5-micron paraffin sections were de-waxed and rehydrated following standard protocols. Antigen retrieval consisted of steaming for 40 min in Target Retrieval Solution (S1700, Agilent, Santa Clara, CA). Slides were then washed and equilibrated in TBS-Tween buffer (Sigma, St. Louis, MO) for 10 min. Primary antibodies were applied at a dilution of 1:25 at 37C for 60 min. For chromogenic studies, immunocomplexes were visualized by applying secondary detection reagents of the UltraVision™ Quanto Detection System (cat. no. TL-060-QHD, Thermo Fisher, Waltham, MA) following manufacturer instructions. Sequential dual-immunofluorescence labeling studies were carried out using Tyramide SuperBoost kits (Thermo Fisher, Waltham, MA). All bright field slides were imaged using a Ventana DP200 system (Roche Diagnostics, Indianapolis, IN). Fluorescence images were acquired on a Cytation 5 Cell Imager (Biotek, Winooski, VT).

### Computational and statistical analysis

#### Analysis of ATAC-seq and ChIP-seq data

A modified version of the ChiLin pipeline was used for quality control and pre-processing of the data^54,55^. We used Burrows-Wheeler Aligner (BWA Version: 0.7.17-r1188) as a read mapping tool to align to hg19 using default parameters. Unique reads for a position for peak calling were used to reduce false positive peaks, and statistically significant peaks were finally selected by calculating a false discovery rate (FDR) of reported peaks. ATAC peaks were called using MACS2 (v2.1.2) with a cut-off of FDR<0.01. H3K27ac, ASCL1 and NEUROD1 peaks were called using MACS2 using the same cut-off. DESeq2 was used to identify differential peaks in ATAC-seq and ChIP-seq, where gained or lost peaks were defined with the threshold of log2 fold change of 1 or 2 and an adjusted p-value<0.05. PCA was performed using princomp in R.

CEAS analysis is used to annotate resulting peaks with genome features. Cistrome Toolkit (dbtoolkit.cistrome.org) was used to probe which factors might regulate the user-defined genes. Genomic Regions Enrichment of Annotations Tool (GREAT) was used to annotate peaks with their biological functions. Conservation plots were obtained with the Conservation Plot (version 1.0.0) tool available in Cistrome^54,55^

#### Analysis of Super-enhancers

Bed files for H3K27ac peaks created by MACS2 were used as input to by ROSE^54^ to call Super-enhancers (SEs) in H3K27ac ChIP-seq data.

#### Visualization of ChIP-seq and ATAC-seq data

Read depth normalized profiles corresponding to read coverage per 1 million reads were used for heatmaps and for visualization using the integrative genomics viewer (IGV)^56^. Heat maps were prepared using deepTools (version 2.5.4) and aggregation plots for ChIP-seq signals were generated using Sitepro in CEAS^57^. In the volcano plots, ATAC-seq peak summits were associated with the nearest TSS within a distance of +/− 50 kb, and incorporating DESeq2 output from RNA-seq, with the final plot generated using ggplot2 in R.

#### Analysis and Visualization of RNA-seq data

For RNA-seq data, read alignment, quality control and data analysis were performed using VIPER^58^. RNA-seq reads were mapped by STAR^59^ to hg19 and read counts for each gene were generated by Cufflinks. Differential gene expression analyses were performed on absolute gene counts for RNA-seq data using DESeq2.

#### Single cell ATAC-seq and RNA-seq

Single cell RNA-seq data generated by 10x Genomics were preprocessed using the Cell Ranger (https://www.10xgenomics.com/) to obtain the UMI (unique molecular identifier) counts for each gene. To get a reliable single cell transcriptome dataset, we excluded the cells with less than 200 genes expressed (UMI > 0) or the cells with more than 80% UMIs from mitochondrial genes. The filtered data was then normalized and scaled by using Seurat^60^ to remove unwanted sources of variations^61^. t-SNE was performed on the normalized data to visualize the single cells in two-dimensional space by using the top 10 dimensions of principal component analysis (PCA). Unsupervised clustering was performed by using the “FindClusters” function in the Seurat package with parameter of resolution = 0.8. Cell cycle phases of all single cells were assigned by using the cyclone function in scran package ^61^. Genes with differential expression between clusters were obtained by using Wilcoxon rank-sum test. FDR was then calculated to correct for multiple testing.

Single-cell ATAC-seq data were processed using the Cell Ranger ATAC pipeline v1.1.0, which provides QC and clustering. Any cell that had Fraction of reads in peaks (FRiP) < 0.2 or total fragments < 1,000 was removed from the analysis. The t-SNE analysis was performed using the implementation from the Loupe Cell Browser 3.1.0.

scATAC-seq and scRNA-seq data integration was performed by Seurat. The scATAC-seq peak matrix provided by 10x was loaded and collapsed to a “gene activity matrix”. The processed data was then scaled and normalized. To help understand the internal structure of the ATAC-seq data, the “RunLSI” function was run. “FindTransferAnchors” function identified ‘anchors’ between the ATAC-seq and RNA-seq datasets, and finally ATAC-seq and RNA-seq data are able to be co-embedded in the same tSNE plot.

### scCNV

By modifying an existing method used for bulk ATAC-seq data, we created a way to use off target scATAC-seq reads to infer DNA copy number amplifications. This approach first breaks the genome into many large intervals and finds the coverage of each window. The coverage of one hundred GC-matched intervals are then averaged together as background. The coverage of each interval will be compared to each GC matched background to estimate CNV fold change. The size of each interval was set to 1-2 Mb to account for the sparsity of the scATAC-seq data with ‘ChunkGRanges’ function in GenomicRange. For each window, the ‘GCcontent’ function of biovizBase was used to calculate the percentage GC content. The coverage was compensated for removed peaks by using the effective window size in coverage calculation.

### Whole exome sequencing

Reads were aligned using BWA v0.5.9 and somatic mutations called using a customized version of the Getz Lab CGA WES Characterization pipeline (https://portal.firecloud.org/#methods/getzlab/CGA_WES_Characterization_Pipeline_v0.1_Dec2018/). We used ContEst^62^ to estimate cross sample contamination, MuTect^63^ v1.1.6 to call single nucleotide variants, and Strelka^64^ v1.0.11 to call indels. MuTect2.1^65^ was used to confirm Strelka indel calls. We applied DeTiN^66^ to rescue true somatic variants that were removed due to tumor-in-normal contamination. Variant calls were filtered through a panel of normal samples to remove artifacts from miscalled germline alterations and other rare error modes. Variants were annotated using VEP, Oncotator, and vcf2maf v1.6.17 (https://github.com/mskcc/vcf2maf). Allelic copy number, tumor purity and ploidy were analyzed using ABSOLUTE^67^.

Prior to characterizing somatic mutations and copy number profiles from patient derived xenograft samples, we removed potentially confounding mouse DNA sequences using ConcatRef^68^. Briefly, WES results were aligned to a concatenated hg19 and mm9 reference genome, and only reads for which both pairs uniquely aligned to just the hg19 reference sequences using BWA. The resultant high-confidence human paired end sequences were then used for downstream analysis as above.

---

**Supplemental Figure 1.**
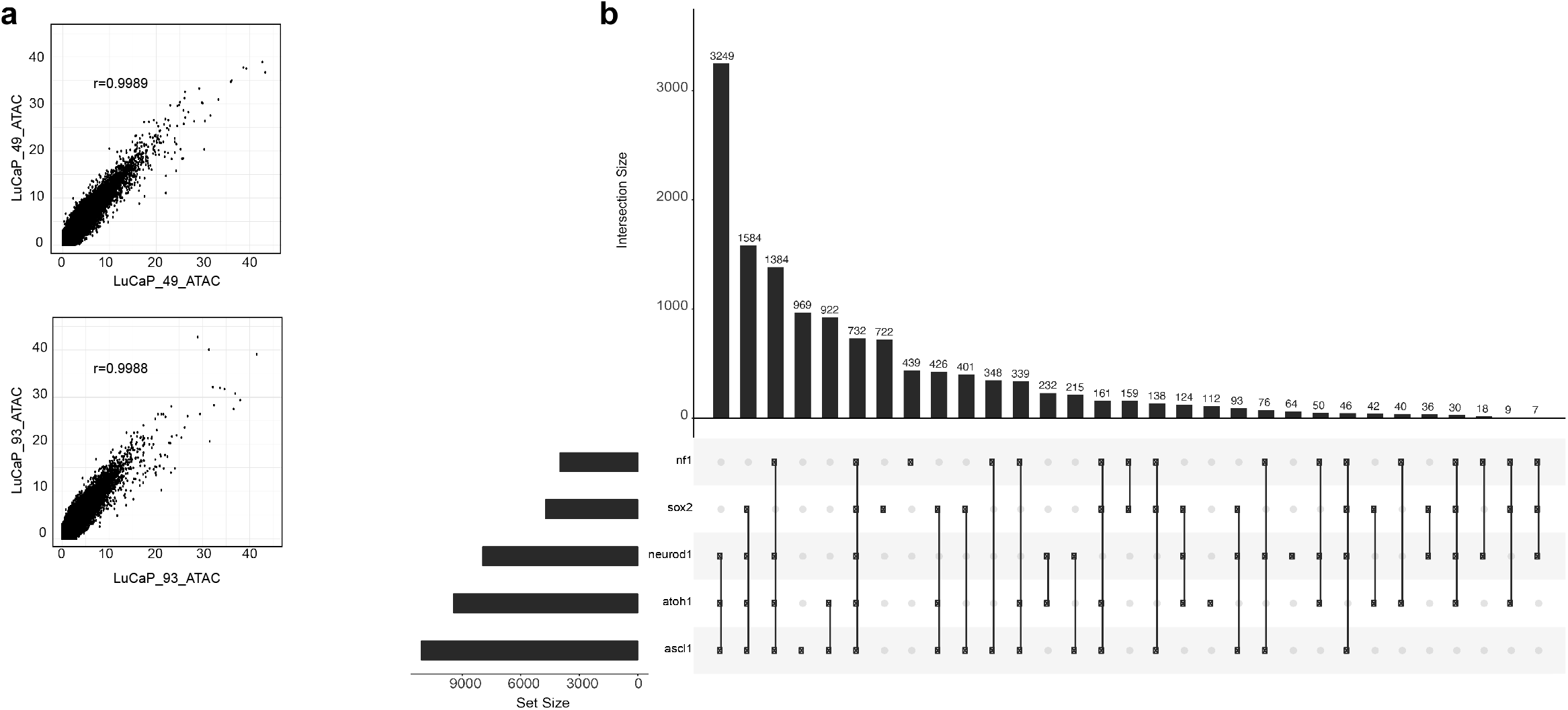
**(a)** Correlation plots of the ATAC-seq signal for two LuCaP_49 replicates (Pearson’s correlation r = 0.9989, top) and LuCaP_93 replicates (Pearson’s correlation r = 0.9988, bottom). **(b)** UpSetR plot^51^ showing the intersections of ATAC-seq peaks containing the specified motifs and their Intersection sizes.

**Supplemental Figure 2.**
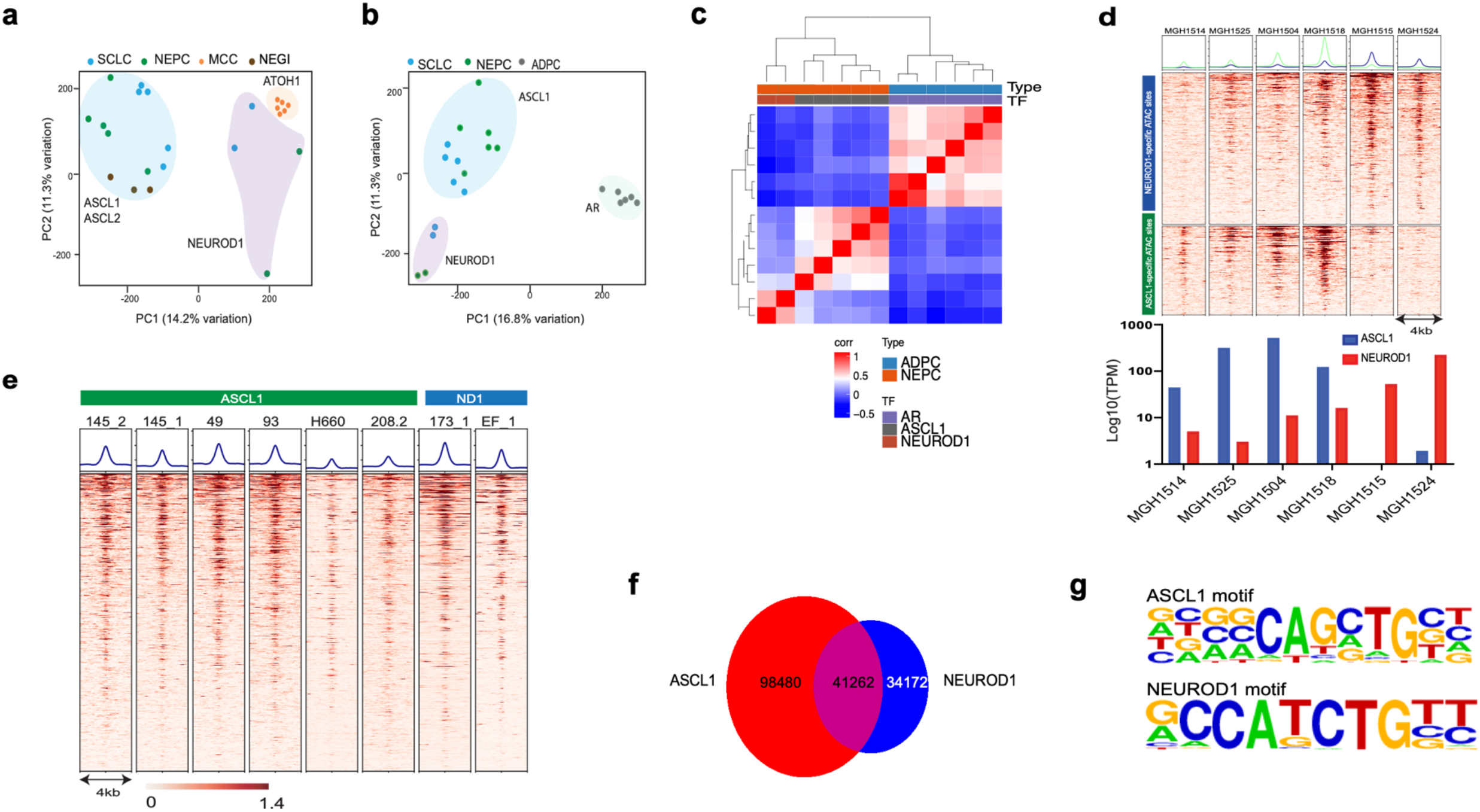
**(a)** PCA analysis of ATAC-seq data of NECs, samples are color coded by tumor type and clusters are highlighted based on the expression of the dominant bHLH TF in each sample. (**b**) PCA analysis of ATAC-seq data of SCLC, NEPC and ADPC, samples are color coded by tumor type and clusters are highlighted based on the expression of the dominant bHLH TF in each sample. (**c**) Hierarchical clustering of the pairwise Pearson’s correlation of the RNA-seq signal across the NEPCs and ADPC. (**d**) Heatmap of the ATAC-seq signal in SCLCs at the ASCL1- and NEUROD1-specific DNA accessible regions identified in NEPC (on top) and corresponding expression of *ASCL1* and *NEUROD1* of the same SCLC tissues (bottom). (**e**) Heatmap of the ATAC-seq signal at the shared accessible regions in the ASCL1 and NEUROD1 NEPC subtypes. (**f**) Venn diagram representing the union of the ASCL1 binding sites obtained by ChIP-seq in PDXs (145.1, 93, 43 and EF1) in red and the NEUROD1 bindings by ChIP-seq analysis in PDX 173.1 (in blue) (**g**) Top enriched consensus motifs identified in ASCL1 ChIP-seq (top) and NEUROD1 ChIP-seq data (bottom).

**Supplemental Figure 3.**
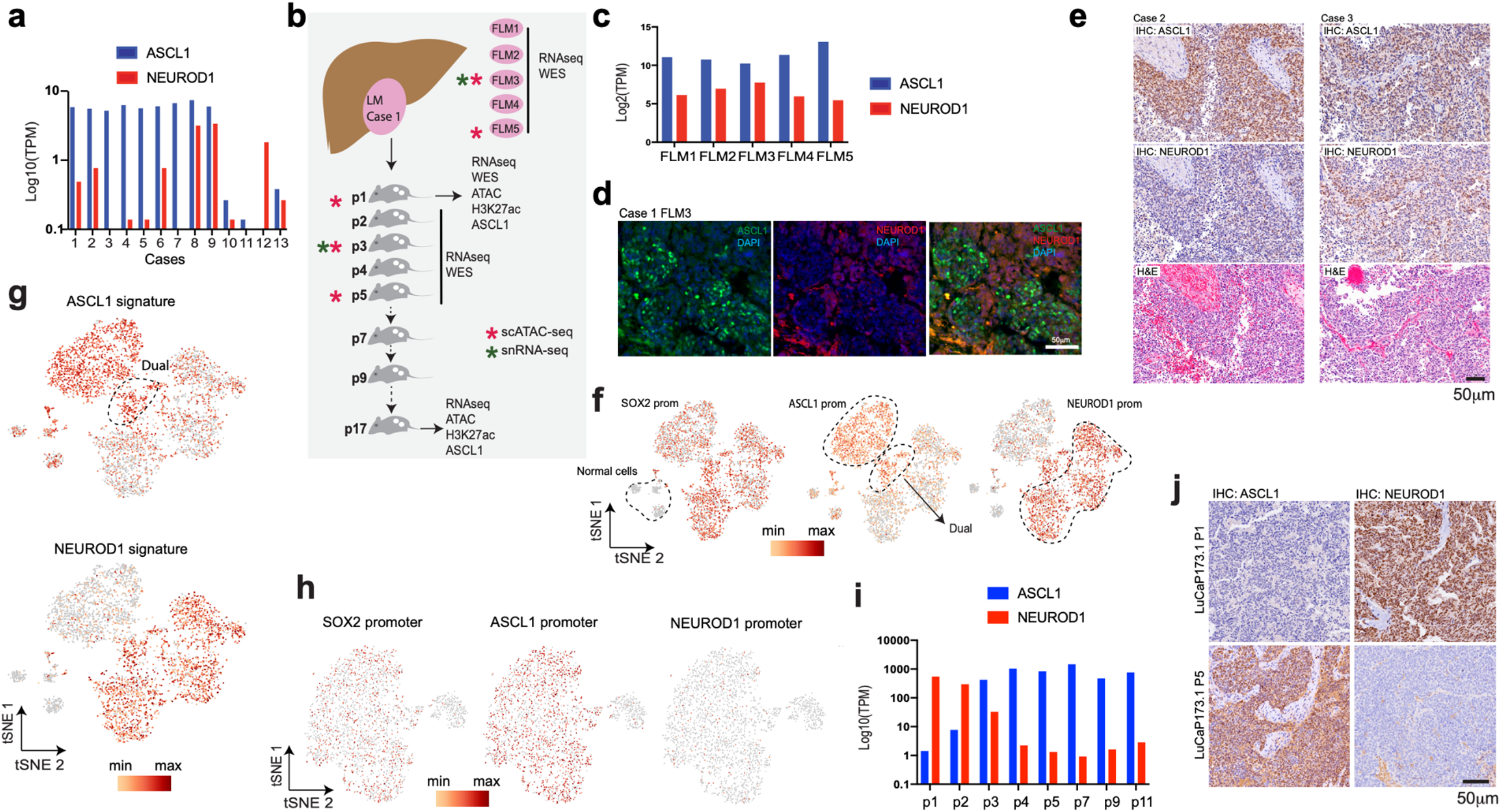
**(a)** Plot of *ASCL1* and *NEUROD1* expression in NEPC tissues from a clinical cohort (Beltran *et al*. 2016)^22^. (**b**) Schematic of the tissues (liver metastasis and tumor passages) analyzed in this study. (**c**) Plot of *ASCL1* and *NEUROD1* expression in the five fragments of liver metastases identified in the case studied (case 1). (**d**) Double immunofluorescence for FLM3 (case 1) with ASCL1 (green) and NEUROD1 (red). (**e**) Immunohistochemical analysis of two additional NEPC liver metastasis cases, ASCL1 on top, NEUROD1 middle panel and H&E bottom. (**f**) t-SNE analysis of the scATAC-seq data of FLM3, showing accessibility at the *SOX2* promoter (marking tumor cells) and differential analysis at *ASCL1* (left) and *NEUROD1* (right) promoters marking ASCL1, “dual” and NEUROD1 subpopulations respectively. (**g**) t-SNE analysis of the scATAC-seq data of FLM3 showing accessibility at the top 30 differential ATAC-seq regions in ASCL1 (top) and NEUROD1 (bottom) subtypes identified by bulk analysis. (**h**) t-SNE analysis of the scATAC-seq data of FLM5, showing accessibility at the *SOX2* promoter (marking tumor cells) and differential analysis at *NEUROD1* (left) and *ASCL1* (right) promoters marking NEUROD1 and ASCL1 subpopulations respectively. (**i**) Analysis of *ASCL1* and *NEUROD1* expression across PDXs passages. (**j**) Immunohistochemical analysis for ASCL1 (left) and NEUROD1 (right) of PDX p1 and p5 showing anticorrelated expression of NEUROD1 and ASCL1 respectively.

**Supplemental Figure 4.**
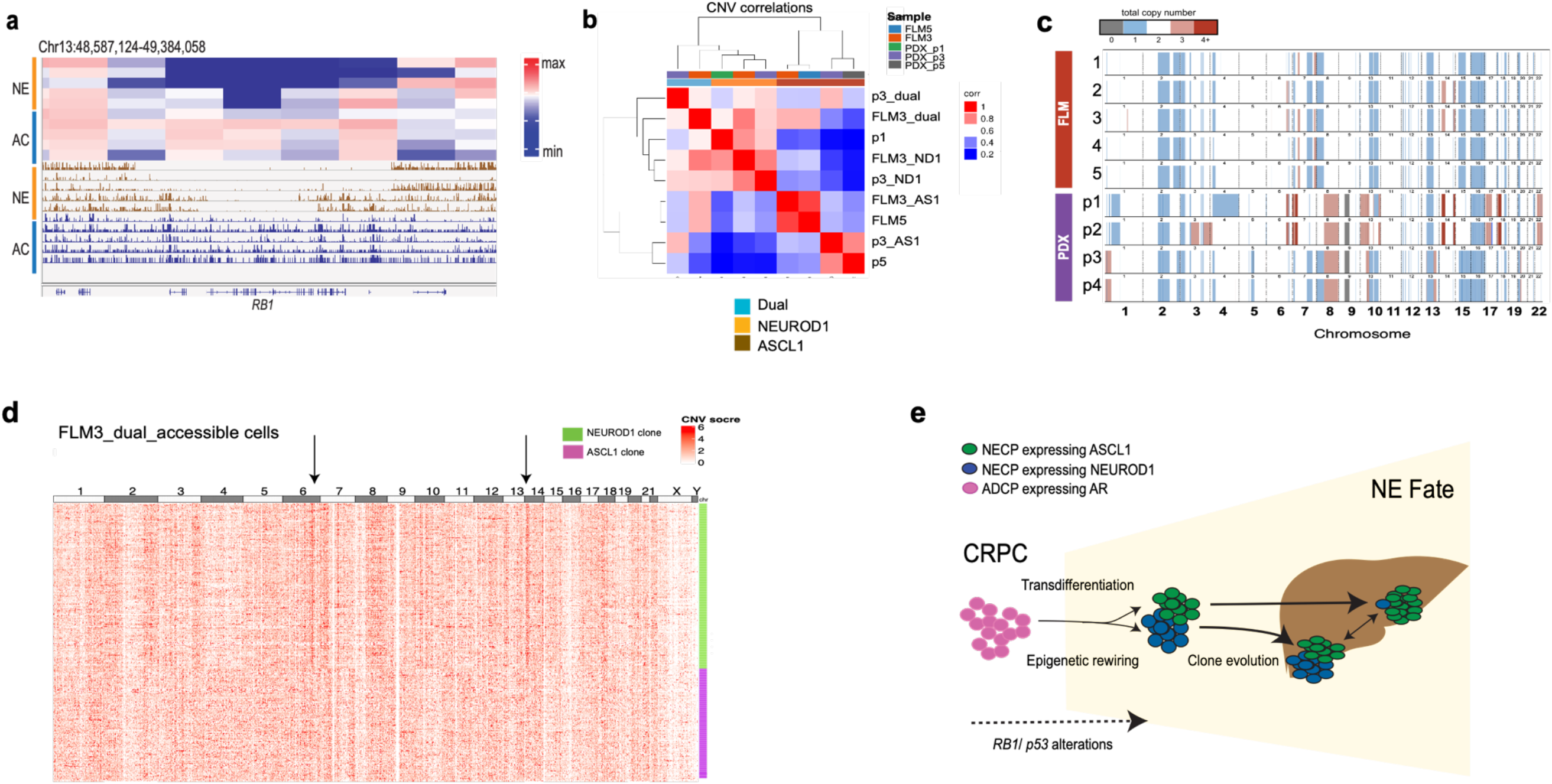
**(a)** Inference of CNV at *RB1* locus from ATAC-seq bulk signal in NEPC and ADPC. (top) Heatmap representation of the inferred data and (bottom) IGV track of the actual ATAC-seq data at that locus. (**b**) Hierarchical clustering of the pairwise Pearson’s correlation of the CNV inference from scATAC-seq t-SNE clusters in FLMs and PDX passages from the case represented in Figure 3b. (**c**) Heatmap of the CNV alterations determined by WES across FLM3 (by clusters), FLM5 and PDX passages (by clusters). (**d**) Heatmap of the single cell CNV analysis of cells showing “dual” accessibility to *ASCL1* and *NEUROD1* promoters. Two dominant clusters are evident that are the same as the larger clusters of ASCL1 and NEUROD1 subclones. (**e**) Model of the origin of the two NEPC subtypes.

**Supplemental Table 1.** Tumor samples used for the ATAC-seq analysis

| Supplemental Table1. Tumor samples used for the ATAC-seq analysis |  |  | ATAC-seq |  |  |  | RNA-seq |
| --- | --- | --- | --- | --- | --- | --- | --- |
| Sample | Tumor | Tissue | UniquelyMapped | Total_Peaks | DHS | FRIP |  |
| DFMC_11112 | MCC | PDX | 45497753 | 91671 | 4897 | 36.4 | Yes |
| DFMC_33043 | MCC | PDX | 50628567 | 90443 | 4828 | 41.1 | Yes |
| DFMC_48396 | MCC | PDX | 56332609 | 94578 | 4871 | 34.8 | Yes |
| DFMC_63632 | MCC | PDX | 44959450 | 66762 | 4671 | 26.7 | Yes |
| DFMC_87346 | MCC | PDX | 47845776 | 61516 | 4887 | 38.6 | Yes |
| DFMC_96712 | MCC | PDX | 44559386 | 72993 | 4832 | 29.4 | Yes |
| EF1_cfce | NEPC | Cell line | 95546177 | 76884 | 4890 | 39.1 |  |
| LuCaP_145_1_rep1 | NEPC | PDX | 27578828 | 25268 | 4806 | 12.6 |  |
| LuCaP_145_1_rep2 | NEPC | PDX | 40326914 | 41537 | 4819 | 4.6 |  |
| LuCaP_145_2 | NEPC | PDX | 21372235 | 32372 | 4892 | 16.6 | Yes |
| LuCaP_173_1_p17 | NEPC | PDX | 37204748 | 31379 | 4828 | 33.3 | Yes |
| LuCaP_173_1_p1 | NEPC | PDX | 173962297 | 79631 | 4782 | 8 | Yes |
| LuCaP_208 | NEPC | PDX | 53527713 | 36861 | 4859 | 9 |  |
| LuCaP_49_rep1 | NEPC | PDX | 40159074 | 35772 | 4881 | 15 | Yes |
| LuCaP_49_rep2 | NEPC | PDX | 44956639 | 51883 | 4893 | 21 |  |
| LuCaP_23 | ADPC | PDX | 30997298 | 49662 | 4868 | 24 | Yes |
| LuCaP_77 | ADPC | PDX | 19527592 | 54493 | 4731 | 26.3 | Yes |
| LuCaP_77CR | ADPC | PDX | 52410587 | 56397 | 4838 | 20.4 |  |
| LuCaP_78 | ADPC | PDX | 29173802 | 48785 | 4641 | 17.6 | Yes |
| LuCaP_81 | ADPC | PDX | 47086721 | 40311 | 4784 | 13.5 | Yes |
| LuCaP_93_rep1 | ADPC | PDX | 39109506 | 35158 | 4664 | 10.5 | Yes |
| LuCaP_93_rep2 | ADPC | PDX | 33565564 | 42470 | 4638 | 14.2 |  |
| LuCaP_96 | ADPC | PDX | 66307812 | 51946 | 4926 | 18.1 | Yes |
| MGH_1504 | SCLC | PDX | 37791305 | 35413 | 4628 | 13.6 | Yes |
| MGH_1514 | SCLC | PDX | 62263801 | 37249 | 4649 | 8.9 | Yes |
| MGH_1515 | SCLC | PDX | 35865754 | 32243 | 4465 | 11.1 | Yes |
| MGH_1518 | SCLC | PDX | 33289869 | 38847 | 4472 | 15.6 | Yes |
| MGH_1524-1 | SCLC | PDX | 45397760 | 25676 | 4704 | 5.7 | Yes |
| MGH_1525 | SCLC | PDX | 42628509 | 27972 | 4758 | 6.8 | Yes |
| MGH_1545 | SCLC | PDX | 35339906 | 51209 | 4697 | 22 | Yes |
| MGH_1567-1 | SCLC | PDX | 58821999 | 21845 | 4910 | 5.8 |  |
| NCI-H660 | NEPC | Cell line | 310339362 | 70104 | 4423 | 15.1 |  |
| T17-11572_NEC | GINEC | Primary tissue | 62189628 | 54148 | 4959 | 22.6 | Yes |
| T18-01479_NEC | GINEC | Primary tissue | 52641558 | 80490 | 4853 | 30 | Yes |
| T18-05426_I_NEC | GINEC | Primary tissue | 58703600 | 55989 | 4896 | 36.2 | Yes |
| DHS: DNase I hypersensitive site (per 5000 peaks) |  |  |  |  |  |  |  |
| FRIP: fraction of reads that fall into a peak |  |  |  |  |  |  |  |

**Supplemental table 2.**
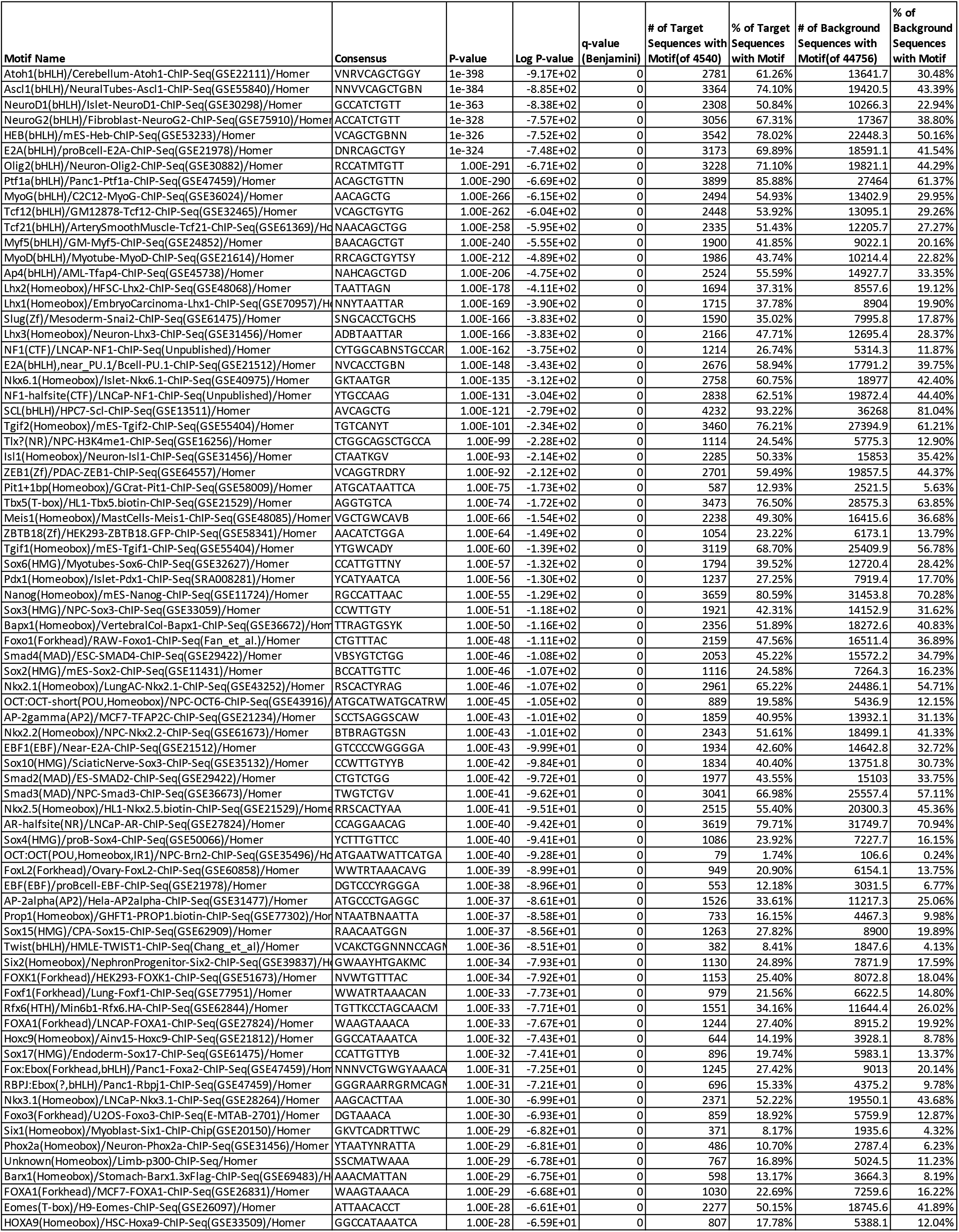
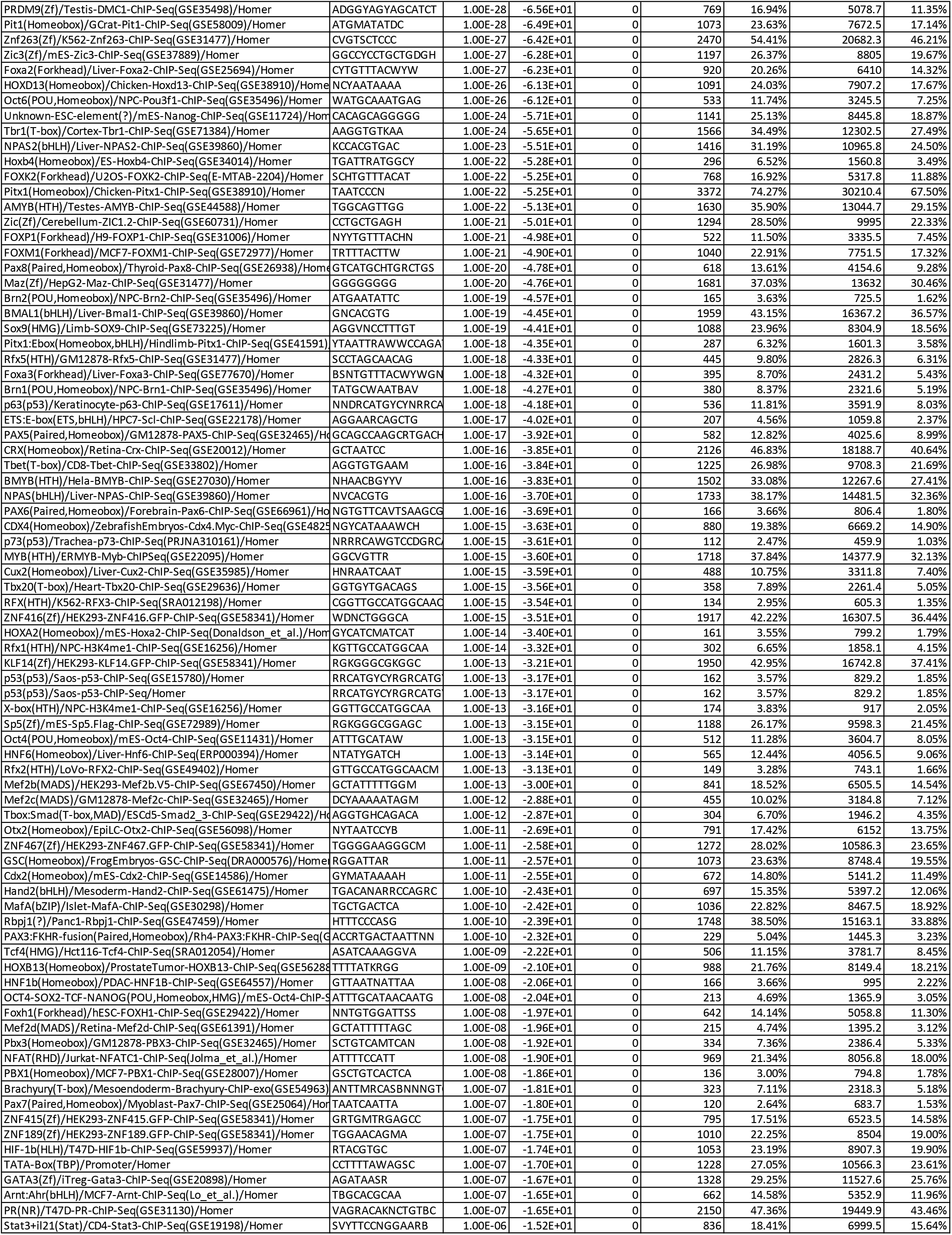
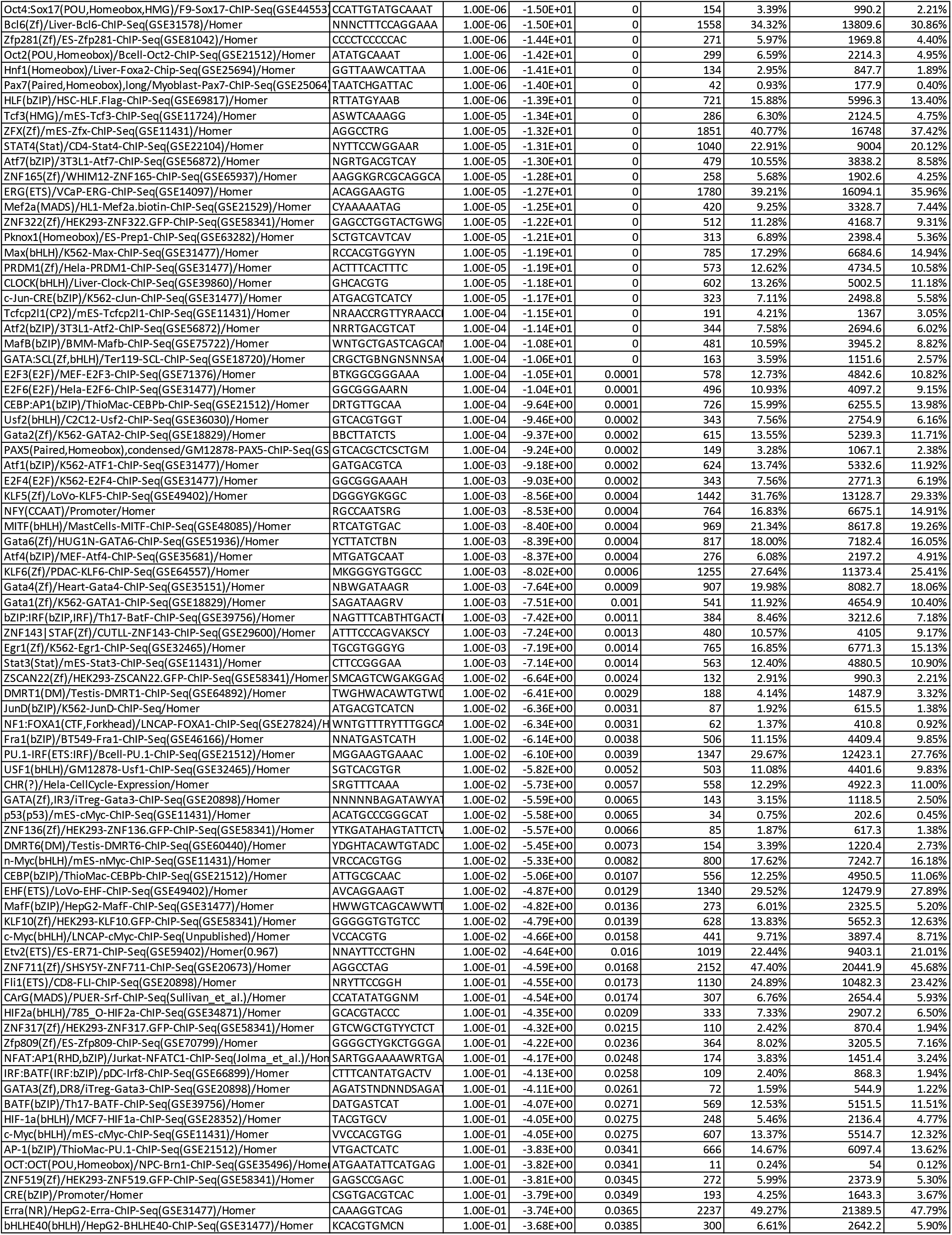
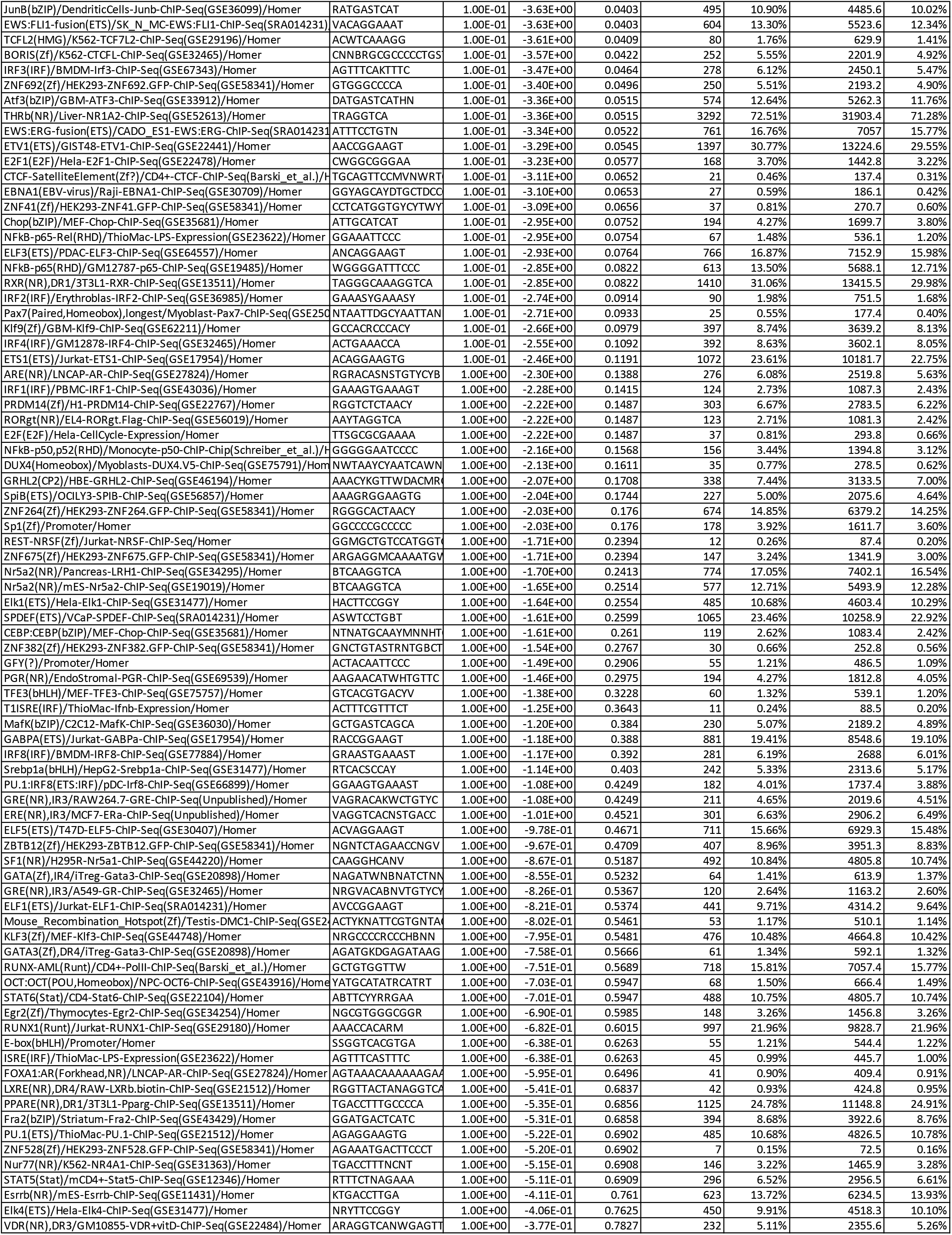
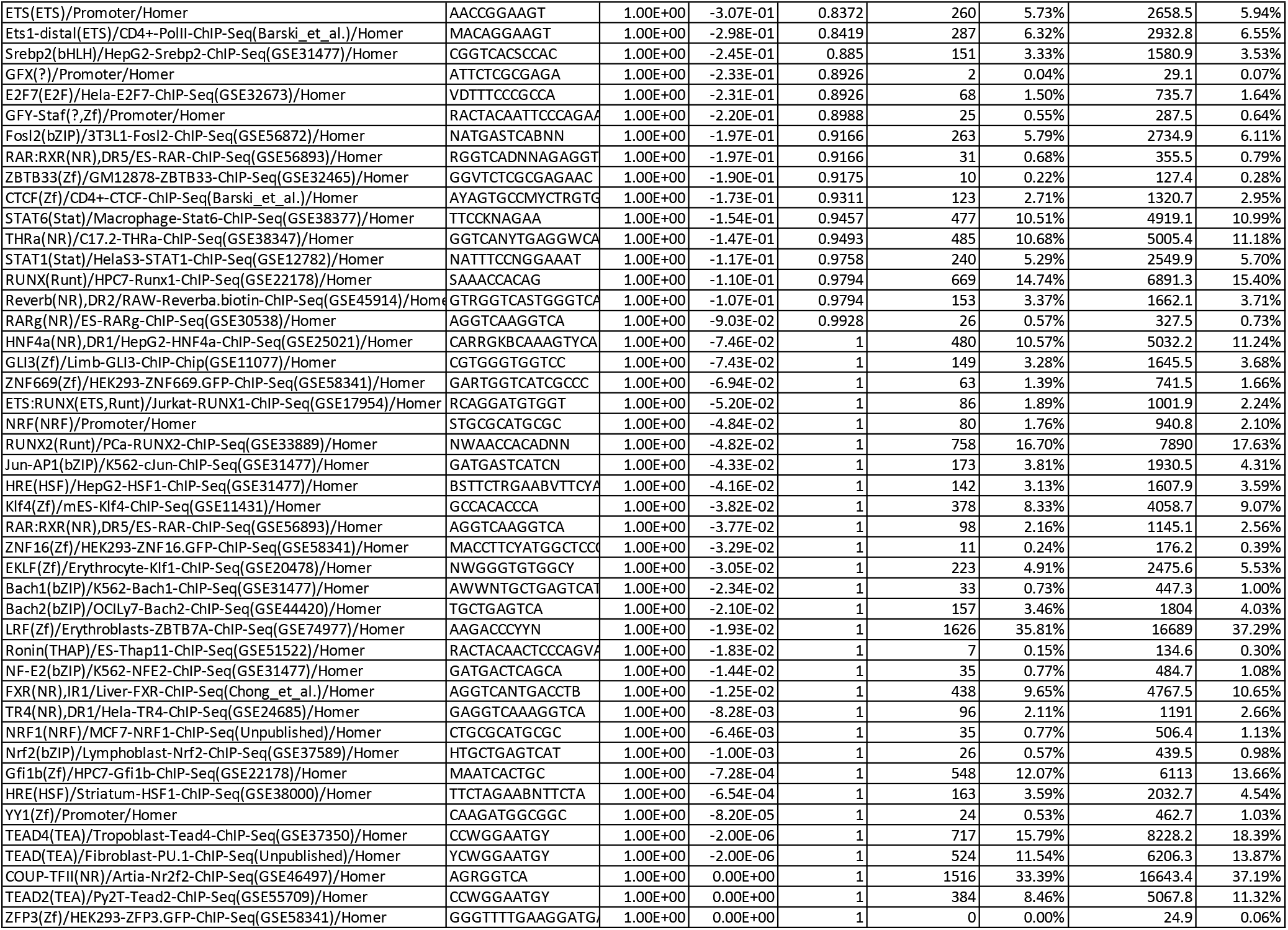
Motif analysis at NE-specific peaks

**Supplemental table 3.** Characteristics of the ASCL1 and NEUROD1 ChIP-seq datasets

| Sample | ChIP-seq | Uniquely Mapped Reads | Total_Peaks | DHS (per 5000 peaks) | FRiP |
| --- | --- | --- | --- | --- | --- |
| LuCaP49 | ASCL1 | 26093095 | 46859 | 4732 | 32.5 |
| LuCaP93 | ASCL1 | 25852594 | 49994 | 4664 | 21.1 |
| LuCaP145_1 | ASCL1 | 38818608 | 46303 | 4259 | 38.9 |
| H660 | ASCL1 | 24979578 | 61338 | 4869 | 34.4 |
| LuCaP173_p1 | NEUROD1 | 31869709 | 78124 | 4857 | 35.1 |

DHS: DNase I hypersensitive site (per 5000 peaks)
FRiP: fraction of reads that fall into a peak

